# Structure guided engineering of a cold active esterase expands substrate range though a stabilisation mutation that allows access to a buried water chamber

**DOI:** 10.1101/2021.02.23.432567

**Authors:** Nehad Noby, Rachel L. Johnson, Jonathan D. Tyzack, Amira M. Embaby, Hesham Saeed, Ahmed Hussein, Sherine N. Khattab, Pierre J. Rizkallah, D. Dafydd Jones

## Abstract

Cold active esterases represent an important class of enzymes capable of undertaking useful chemical transformations at low temperatures. EstN7 from *Bacillus cohnii* represents a true psychrophilic esterase with a temperature optimum below 20°C. We have recently determined the structure of EstN7 and have used this knowledge to understand substrate specificity and expands its substrate range through protein engineering. Substrate range is determined by a plug at the end of acyl binding pocket that blocks access to a buried water filled cavity, so limiting EstN7 to turnover of C2 and C4 substrates. Data mining revealed a potentially important commercial reaction, conversion of triacetin to only the 1,2-glyceryl diacetate isomer, which the EstN7 could achieve. Residues M187, N211 and W206 were identified as plug residues. M187 was identified as the key plug residue but mutation to alanine destabilised the structure as whole. Another plug mutation, N211A had a stabilising effect on EstN7 and suppressed the destabilising M187A mutation. The M187A-N211A variant had the broadest substrate range, capable of hydrolysing a C8 substrate. Thus, the structure of EstN7 together with focused engineering has provided new insights into the structural stability and substrate specificity that allowed expansion of substrate range.

## Introduction

The urgent need for sustainable eco-friendly chemical approaches has led to the search for alternatives to traditional catalysts (1,2). Enzymes are one such source due to their functional plasticity, need for less harsh conditions (low temperature and pressure, low organic solvent use) and ability to be biodegraded. (3–5). Esterases are especially useful as they can utilise a variety of ester substrates, catalysing their hydrolysis into their constituent alcohol and acid (6–8). Cold active/adapted enzymes are attracting particular attention as their optimal activity at lower temperatures can be applied to temperature sensitive processes (9–14) such as: the pharmaceutical industry for synthesising fragile chiral compounds (15,16); detergent industry for cold washing (17); environmental bioremediation (18). Cost competitiveness is a fundamental consideration for implementing bio-catalysis in the industry. Usually, the environmental advantages alone cannot support the replacement of conventional chemical catalysts. Enzymes with high activity, required specificity and stability are key determinants to guarantee high performance and low-cost production approaches. However, the catalytic properties of native enzymes may not fully meet the industrial requirements leading to them being engineered to better fulfil a particular requirement (19).

We have recently determined the structure of cold active esterase, termed EstN7, from *B. cohnii* strain N1 (20) (Figure 1a). Unlike many esterases isolated from psychrophilic organism that still have optimal activity at >20°C, EstN7 is truly cold active with optimal temperature <20°C (21). Its low temperature activity profile is coupled to relatively high thermal stability, with a melting temperature of 51°C. EstN7 also retained function in the presence of up to 30% of various organic solvents, which makes it potentially useful in various applications, such as fine chemical synthesis and pharmaceutical industries. The natural function of the enzyme is currently unknown and little was known about its structure until recently (20). Analysis of the structure and dynamics of EstN7 suggests a water entropy-based cold adaptation mechanism rather than changes in local and global dynamics (20); the surface of EstN7 is optimised to make interactions with water and so retaining the water shell even at lower temperatures when water viscosity increases. The retention of a water shell may also explain EstN7’s tolerance to organic solvents (12). We also speculated that part of the cold adaption mechanism is improved access to the active site. The N-terminal cap (Figure 1a) is common to esterases and plays a critical role in function (7,13); it contributes to forming the walls of the main channel by which substrate accesses the active site, which is located at the bottom of a cleft. In EstN7, the N-terminal cap is loosely associated with the main catalytic domain and forms a “bridge” like structure which results in additional channels for the substrate to reach the active site (Figure 1b). These additional channels are missing from the closest related mesophilic and thermophilic structural homologues (20).

**Figure 1.**
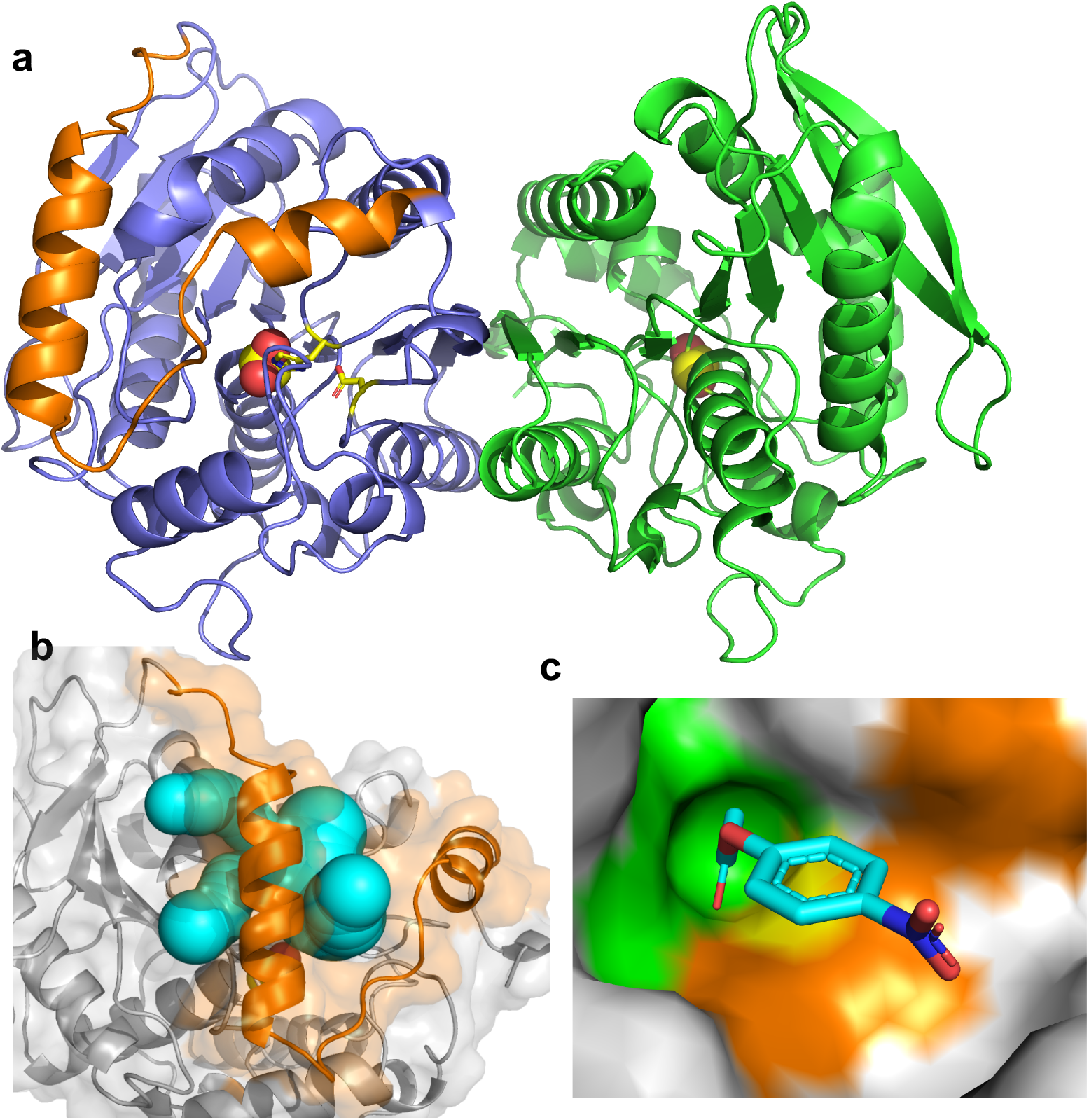
Structure of EstN7^WT^. (a) Overall dimeric structure of EstN7 (subunit A and B coloured blue and green, respectively) with the catalytic residues coloured yellow. Details of the catalytic site are shown in Figure S1a. (b) Channels available for substrate to enter the active site (20). Channels are coloured in cyan and were identified using CAVER 3.0 (22) with a minimum probe radius of 1.4 Å and maximum distance of 10 Å. (c) Substrate binding regions with *p*-NP-C2 docked. The green region represents the acyl binding site, orange the alcohol binding site and yellow the catalytic residues. A simplified scheme is shown in Figure S1b.

Here we focus on the substrate specificity of EstN7 and how knowledge of its structure guided our engineering efforts to expand substrate range. We find that the specificity of EstN7 is restricted to short chain compounds only. Using a bespoke version of the enzyme biotransformation datamining software Transform-MinER, built on known esterase reactions, we have identified useful putative substrates such as triacetin. Using structure-guided protein engineering we also successfully expanded EstN7’s substrate range from exclusively short carbon chain substrates to longer chain substrates (up to C8). Identification of a suppressor mutation (N211A) that comprised the acyl binding site proved critical to expanding substrate specificity.

## Results

### EstN7^WT^ has a restricted substrate range

The basic physicochemical properties of EstN7 have been reported by Noby et al elsewhere, including optimal activity at 5-20°C, temperature-dependent stability based on activity, organic solvent stability and metal ion sensitivity (21). However, the substrate specificity of EstN7 is currently unknown, as is the case with many natural esterases. We have recently reported the crystal structure of (20) (Figure 1). Briefly, EstN7 is dimeric with a α/β fold similar to that found in other family IV esterases (6,23). The catalytic triad is comprised of S157, H284 and D254 (Figure S1a), and is located at the bottom of a cleft. Two separate rotamer configurations were observed for the Ser157 nucleophile, which are thought to represent actual conformations (24–26); rotamer 1 in Figure S1a represents the catalytically competent state while rotamer 2 is thought to aid with product release so preventing cycling with the reverse reaction. Interestingly, an ethylene glycol moiety was found bound within the active site might that may act as a substrate/product mimic given that one alcohol group is within H-bond distance of S157 hydroxyl group (Figure S1a).

Enzyme kinetics with a range of *p*-nitrophenyl (pNP) substrates with different length carbon chains shows that EstN7 also has a preference for short chain substrates (Table 1). The *p*-NP acetate (C2) is by far the preferred substrate with the *p*-NP butyrate (C4) substrate ~3 orders of magnitude less efficient. There was no detectable activity for longer substrates *p*-NP hexanoate (C6) and *p*-NP octanoate (C8). Comparing the C2 and C4 kinetics, the major change was in turnover, with a ~2300-fold decrease in *k*_cat_ with *K*_M_ remaining similar. This suggests that C2 and C4 bind with similar affinity but that C2 is bound in such a manner that it is more likely to undergo catalysis.

**Table 1.** Substrate specificity of wildtype EstN7and its engineered variants.

| EstN7 variant | p-NF Substrate | $K_M$ (mM) | $k_{cat}$ ($s^{-1}$) | Catalytic efficiency ($s^{-1} M^{-1}$) | Fold change <sup>a</sup> |
| --- | --- | --- | --- | --- | --- |
| WT | C2 | 0.05 ±0.007 | 760 ±3.5 | 1.52x10<sup>7</sup> | 1 |
|  | C4 | 0.057 ±0.002 | 0.32 ±0.005 | 5614 | 1 |
|  | C6 | - | - | - | - |
|  | C8 | - | - | - | - |
| N211A | C2 | 0.75 ±0.001 | 280.5 ±0.07 | 3.74 × 10<sup>5</sup> | 0.024 |
|  | C4 | 2 ±0.2 | 15 ±0.5 | 7500 | 1.33 |
|  | C6 | 0.374 ±0.05 | 2.35 ±0.21 | 6283 | - |
|  | C8 | - | - | - | - |
| W206A | C2 | 1.3 ±0.14 | 1.55 ±0.07 | 1192 | 7.8x10<sup>-5</sup> |
|  | C4 | 1.2 ±0.55 | 8 ±0.98 | 6666 | 1.2 |
|  | C6 | 1.15 ±0.07 | 3.85 ±0.07 | 3347 | - |
|  | C8 | - | - | - | - |
| DM<br>(M187A /<br>N211A) | C2 | 1.1 ±0.07 | 361 ±0.14 | 3.28 × 10<sup>5</sup> | 0.021 |
|  | C4 | 0.3747 ±0.08 | 17.55 ±2.1 | 4.68 × 10<sup>4</sup> | 8.0 |
|  | C6 | 1.2 ±0.01 | 24.5 ±0.7 | 2.04 × 10<sup>4</sup> | - |
|  | C8 | 0.95 ±0.02 | 0.45 ±0.02 | 473 | - |
| TM<br>(M187A /<br>W206A /<br>N211A) | C2 | 2 ±0.3 | 0.07 ±0.012 | 35 | 2.31 x10<sup>-6</sup> |
|  | C4 | 2.1 ±0.36 | 1 ±0.11 | 467 | 0.083 |
|  | C6 | 0.59 ±0.026 | 1.45 ±0.021 | 2457 | - |
|  | C8 | 2 ±0.03 | 0.85 ±0.007 | 425 | - |

a, compared to EstN7<sup>WT</sup> activity on the same substrate.

### Structural basis for substrate specificity

Analysis of the EstN7 structure and substrate docking provide rationale for the restricted substrate range to short chain units. The initial step in substrate binding is access through to the active site, which in family IV esterases is normally through a single channel in which the cap region forms the walls. Previous channel analysis of EstN7 revealed two potential channels through to the active site; a primary channel common to other esterases and secondary channel formed due to the bridge-like structure of the second helical region in the cap domain (Figure 1b).

We docked the primary substrate, pNP-C2, into the active site of EstN7 manually in COOT(28), followed by geometry regularisation with REFMAC(29). In terms of the pNP substrates, the nitrophenyl group comprises the alcohol component and the linear alkane chain the acyl moiety (Figure S1b). The C2 substrate binds to place the hydrolysable ester bond close to Ser157 (Figure 2a). The active site of esterases is comprised of the acyl binding site and the alcohol binding site, outlined in Figure 1c, and form two distinct pockets. The alcohol binding site is open and directly faces the channel opening. The EG^1^ molecule in the active site (Figure S1b) essentially binds within the alcohol binding pocket, in keeping with its chemical character. The acyl binding pocket is buried further into the protein and has a much more restricted volume. It is the acyl binding pocket that restricts the activity of EstN7 to the short chain substrates used here. Residue M187 together with N211 form a plug at the base of the acyl binding pocket so preventing access of longer chained substrates from binding, with M187 providing most of the steric blocking (Figure 2a). Comparison of EstN7 with the closest structural relatives provides further insight into the substrate specificity around the plug (Figure 2b and S2). The closest relative, the putative heroin esterase HerE (PDB 1LZK) (26) has an acidic plug due to replacement of methionine (M187) with a glutamate (E190) (Figure S2a-b). The asparagine (N211) in EstN7 is replaced by an alanine (A214) but this is not enough to fully open the HerE plug. LipW (PDB 3QH4) (25) has two shorter residues in place of M187 and N211 (V192 and A215, respectively) but the plug is completed by M218 that positions the sulphur residue at a position similar to that of M187 (Figure S2c-d). PestE (PDB 2YH2) (27) does not have the plug (Figure 2b). N211 is replaced by a methionine (M214) but the backbone trajectory is away from the plug. M187 is replaced with alanine (A187) and the W206 with a leucine (L209). These sequence and conformational changes remove the plug allowing PestE to hydrolyse longer chain substrates (30).

**Figure 2.**
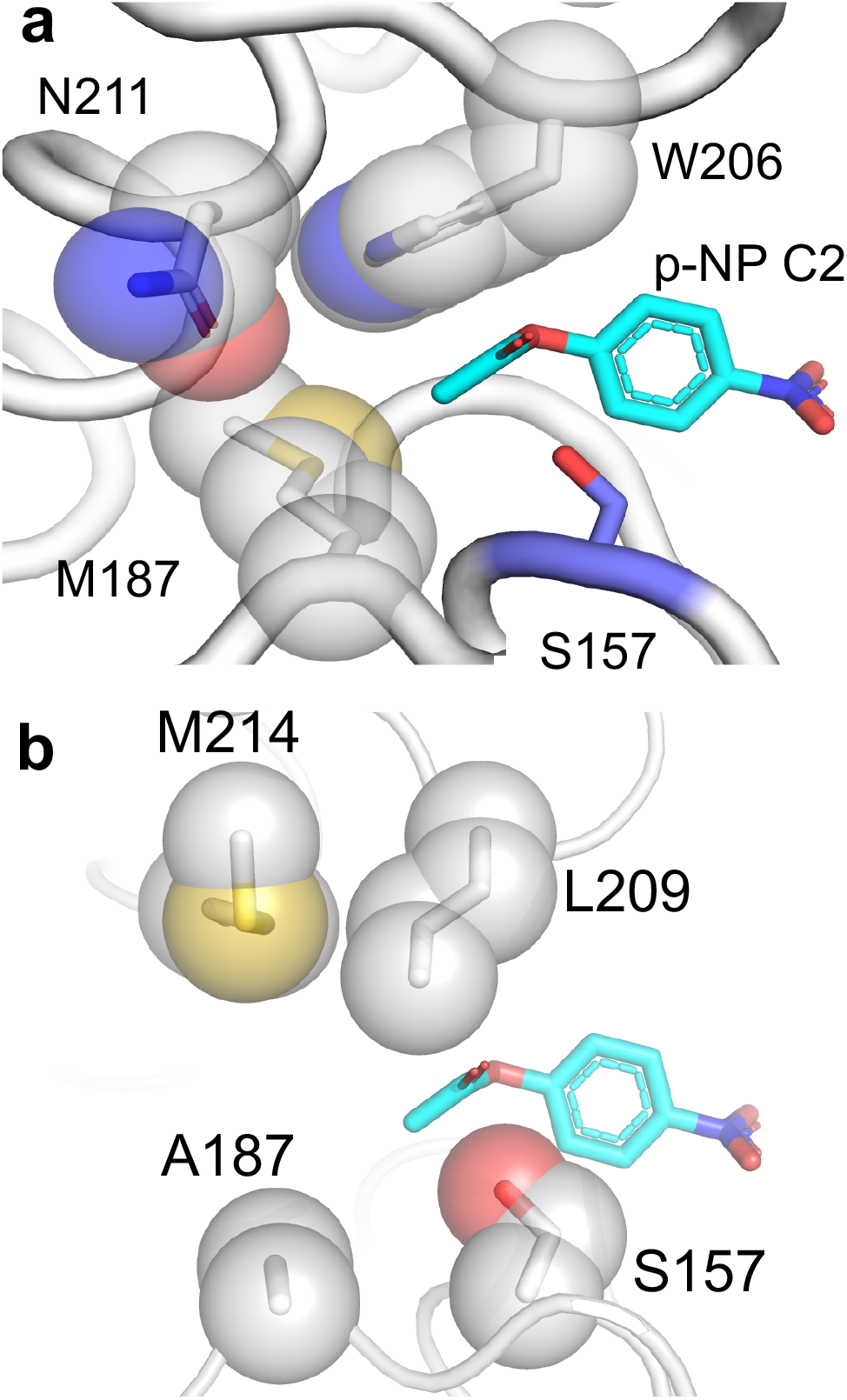
Structural basis for substrate binding and specificity in EstN7. The docked substrate pNP-C2 is coloured cyan. (a) Positioning of three residues contributing to the acyl binding pocket and the plug structure; (b) The open binding acyl binding pocket in PestE (27) (PDB 2YH2). Subunit A is shown for reference.

### Identification of potential substrates for EstN7

Virtual screening based on the EstN7 adapted version of Transform-MinER (31) identified a number of reactions with high similarity to known reactions with potential commercial-value based on recent purchase prices. The majority of these were in the ester forming direction. However, one reaction identified with potential commercial value was hydrolysis of triacetin (glyceryl triacetate) at the terminal acetyl position to form 1,2-glyceryl diacetate (1-(acetyloxy)-3-hydroxypropan-2-yl acetate). Triacetin is the simplest of all the triacylglycerides (Figure 3a); the three alcohol groups of glycerol are acetylated. Triacetin is itself an important commercial compound with its uses including in the pharmaceutical industry as a biodegradable gel system for drug delivery (32). Glyceryl diacetate, normally in the form as an isomeric mixture of the 1,2 and 1,3 forms (Figure 3a), has numerous uses in the industrial and consumer sectors, including the food industry. In our analysis of recent purchase prices, the terminal mono-deacetylated product is highlighted as more valuable: for instance, supply of 10g of the terminal mono-deacetylated product is ~$3,000 compared to 10g of triacetin for $27.

**Figure 3.**
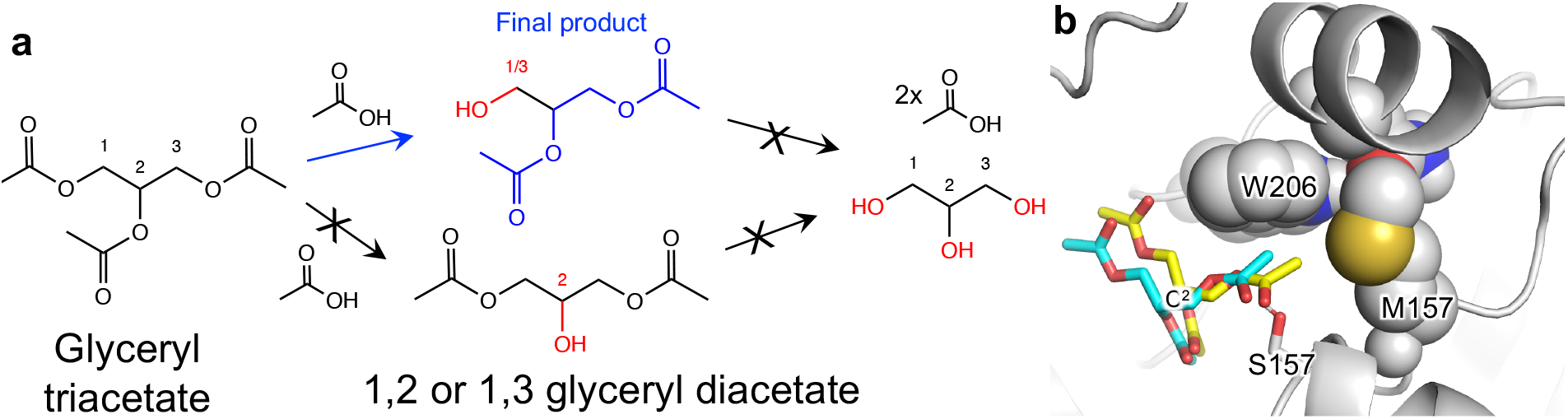
Triacetin as a substrate. (a) Potential routes of triacetin hydrolysis. The final observed product is shown in blue (1,2-glyceryl diacetate) with other potential products (1,3-glyceryl diacetate, the monoacetates or glycerol) not observed coloured black. (b) Triacetin docking to EstN7. Two docked forms of triacetin (yellow and cyan) are shown. The cluster represented the lowest HADDOCK score (−24.0±0.6) observed for the simulation. The central C2 carbon of the glycerol backbone is highlighted for reference.

The key question is whether EstN7 will remove just one of the terminal acetyl groups thus triacetin was tested as a substrate for EstN7. There are three potential hydrolysable ester bonds in triacetin (Figure 3a). Substrate docking of triacetin to EstN7 suggested that one of the terminal ester linkages would be the target for hydrolysis (Figure 3b). Analysis of the reaction products reveals that only one acetyl group is liberated during the course of the EstN7 catalysed reaction, with NMR analysis of the reaction products suggesting either hydrolysis the terminal 1 or 3 position and not in the central C2 carbon position (Figure S3 and accompanying analysis). Thus, EstN7 can potentially be used to convert triacetin into the commercially valuable and isomerically pure 1,2-glceryl diacetate.

### Expanding EstN7 substrate range by structure guided engineering

Based on analysis of the EstN7 structure, three residues considered critical for restricting the enzyme’s specificity to C2 substrates were targeted for mutagenesis: M187, N211 and W206. Behind the plug lies a water-containing cavity (Figure 4a); these water molecules are effectively trapped within the interior of the protein. The presence of a water-filled cavity beyond the plug is also observed in related esterase structures, with the number of waters varying from one (HerE) to three (LipW).

**Figure 4.**
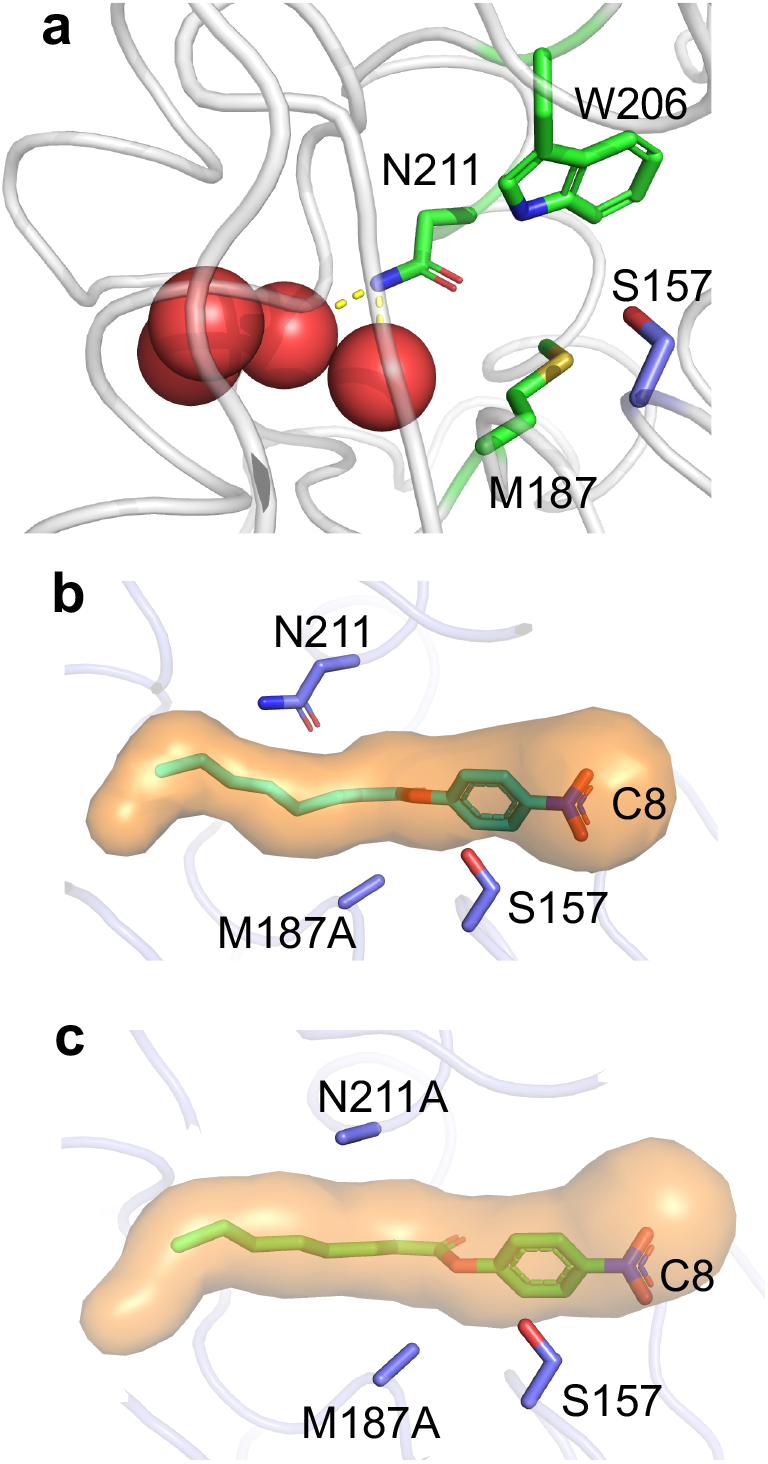
Opening the acyl pocket plug. (a) The cavity behind the plug containing water molecules (red spheres). The nucleophilic S157 is coloured blue and the plug residues M187, W206 and N211 are coloured green. Polar contacts from the carboxamide to the water molecules are shown as yellow dashed. *In silico* modelling of acyl plug mutations and docking of substrate C8 for the (b) M187A mutation alone and (c) in combination with N211A.

*In silico* modelling of the M187A mutation suggests that the plug will open so allowing substrates up to C8 to bind (Figure 4b). However, the M187A mutation had a major disruptive effect on the folding and stability as EstN7^M187A^ was produced in *E. coli* as inclusion bodies. We next targeted N211A. While modelling suggested that N211A would not totally remove the plug, the protein was produced as soluble and functional. Most notable was the effect of N211A on stability. Circular dichroism (CD) spectroscopy was performed at 20°C (Figure 5). Both proteins showed helical signatures, with troughs at 208 nm and 222 nm suggesting that at 20°C, the proteins were folded. EstN7^N211A^ had a higher proportion of helical character than the WT suggesting a higher degree of structure for the mutant at this temperature. Thermal unfolding of EstN7^N211A^ monitored by CD at 222 nm revealed a major transition between 55°C and 65°C, with a T_m_ of 60°C (Figure 5b). As we reported previously (20), EstN7^WT^ has a more extended transition (45°C-60°C) with a shallower slope and a T_M_ of 51°C. Both EstN7^WT^ and EstN7^N211A^ had similar spectra after 70°C suggesting that they had reached the same structural endpoint (i.e. denatured) at the end of temperature ramping (Figure S4).

**Figure 5.**
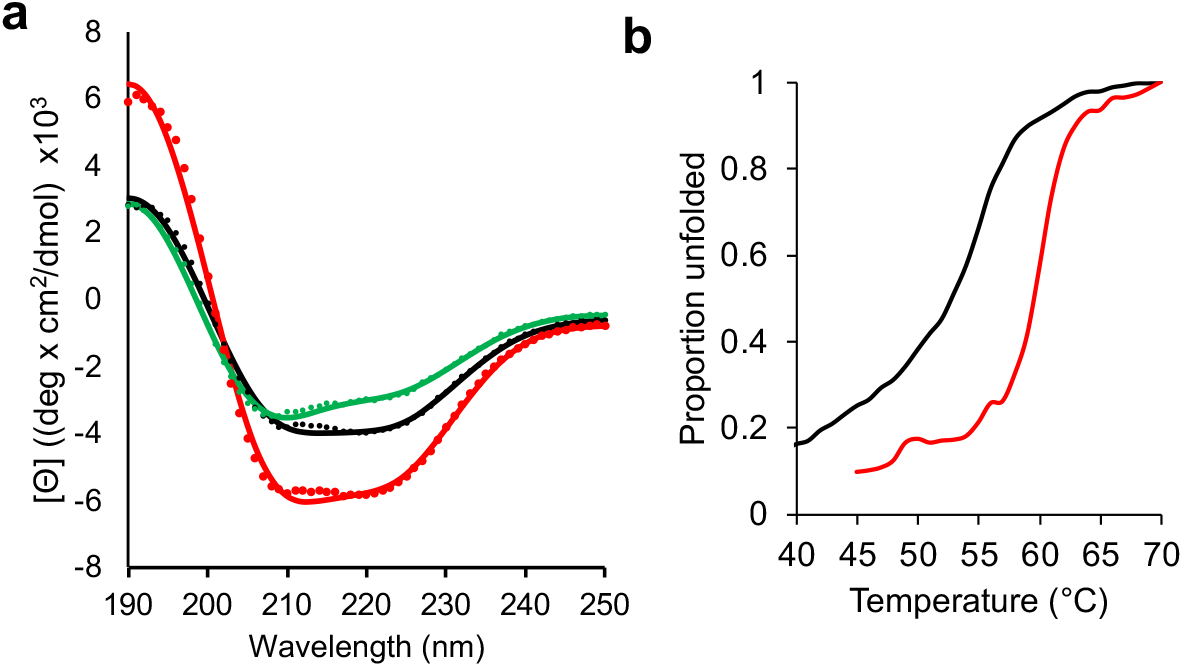
CD spectroscopy of EstN7. (a) CD spectra of EstN7^WT^ (black), EstN7^N211A^ (red) and EstN7^DM^ (green). CD spectra were collected at 20°C and a Savitzky-Golay filter applied. (b) Temperature-dependent changes in molar ellipticity at 222 nm for EstN7^WT^ (black) and EstN7^N211A^ (red).

The N211A mutation improved catalytic efficiency towards C4 substrate by 2.4 fold compared to EstN7^WT^, with *k*_cat_ increasing 30 fold. The consequence was a drop in activity towards C2, with EstN7^N221A^ retaining ~2% of the catalytic efficiency of EstN7^WT^; the biggest contributor was a 15-fold increase in *K*_M_. Importantly, the N211A mutation opened up the ability to hydrolyse C6 but not C8 substrate (Table 1).

As the N211A mutation has a stabilising effect on EstN7, a double mutant which also incorporated the M187A mutation was generated. The double mutant (EstN7^DM^) protein was produced as a soluble and active protein, with the CD spectrum having a similar profile to EstN7^WT^ (Figure 5a). This suggests that the stabilising effect of N211A offset the disruptive nature of the M187A mutation. EstN7^DM^ had observable activity on all tested substrates (C2 to C8) demonstrating that the mutations had expanded substrate range of the esterase (Table 1). Activity towards C2 was still significant but as with EstN7^N211A^ was ~2% that of the wild-type enzyme suggesting a shift in substrate specificity. Activity towards C4 is nearly an order of magnitude higher compared to EstN7^WT^, with *k*_cat_ just over 50-fold higher. EstN7^DM^ activity towards C6 is also improved compared to EstN7^N211A^, with *k*_cat_ increasing almost 10-fold. Importantly, EstN7^DM^ is now active towards C8 substrates.

W206 appears to be important for EstN7 function as mutation to alanine had a major effect on both *k*_cat_ and *K*_M_ with catalytic efficiency for C4 lower than that observed for EstN7^WT^ (Table 1). W206 forms two hydrophobic interactions in chain A and three in chain B. W206A did appear to shift substrate specificity towards longer chained substrates with a 5.4 and 2.7 fold increase in catalytic efficiency to C4 and C6 compared C2 as a substrate. Incorporating W206A into the EstN7^DM^ to create the triple mutant had little overall effect, with catalytic efficiency towards C2, C4 and C6 dropping 5, 85 and 10,000 fold, respectively, compared to the double mutant (Table 1); the triple mutant did display the highest catalytic efficiency towards C8 but only slightly and was largely similar to EstN7^DM^.

## Discussion

The inherent reaction catalysed by esterases is important for various industrial applications but limits on substrate range can impose a major restriction. The physiological function of EstN7 is unknown but given its high activity towards short chain substrates (Table 1) its main role may not be a lipase that hydrolyse long chain acyl units. For example, HerE has a similarly size restricted acyl binding pocket (Figure S1), which may account for its presumed heroin esterase (26). Using substrate mining approaches, we found a potentially useful substrate for EstN7, triacetin (also known as glyceryl triacetate; Figure 3). Ideally, an esterase will be able to catalyse turnover of a range of substrates thus introducing the idea of functional plasticity or at least be catalytically precise so generating a desired single product (13,33), as demonstrated here with triacetin being hydrolysed at only a single terminal acetyl unit (Figure 3a and S3). However, triacetin is a synthetic molecule so is unlikely to be the natural substrate for EstN7.

There has been a lot of focus on identifying natural esterases with the required activity (34,35) or engineering existing esterases to widen their substrate range (36,37). Various studies have shown that altering the residues of the acyl binding tunnel can alter enzyme chain length specificity (38,39). Here, we focused on 3 residues that structural analysis and substrate docking suggests contributes via a structural component we refer to as the “plug” (Figure 1c and 2a), which restricts substrate range to short chained carbon units. The plug is common to some of the close relatives of EstN7 such as Est8 (40), LipW (25) and HerE (26) (Figure S2). In contrast, the thermophillic PestE does not have a plug (Figure 2a), which in turn leads to its ability to hydrolyse longer chained substrates (27,30). Thus, to open the plug M187, W206 and N211 were targeted; based on our modelling, mutating the residues to alanine should open the plug and allow substrates up to C8 to bind (Figure 4b-c).

An unexpected but important observation is the role of the N211A mutation in stabilising EstN7 that allowed expression of the destabilising M187A mutation (Figure 5). Rigidity analysis undertaken previously (20) revealed that M187 forms four hydrophobic interactions, indicating that it is involved in the network of hydrophobic interactions stabilising the protein’s core structure. The loss of so many interactions on mutating M187 to alanine may account for its destabilising effect. N211 is buried and lies in the first turn of helix 7 just above the active site (Figure 2a and 4a). While none of the plug residues (including N211) form strong polar sidechain interactions with other residues, N211 makes polar contacts with two of the four buried water molecules that lie beyond the plug. Additional residues make up the cavity having the potential to form mainly backbone H-bonds to the waters (Figure S5). While related esterases contain water molecules within this cavity, none contain four trapped water molecules; trapping such water molecules may come with a high entropic cost to EstN7 as the temperature rises. While loss of hydrophobic interactions is likely to play a major role regarding the destabilisation effect of M187A, additional local changes on mutation could disrupt the interaction network required to stabilise the internal water molecules. To offset this effect, the N211A mutation may result in loss of a protein-water interaction and generate a potential tunnel for waters to escape so providing an entropic contribution to stabilisation. N221 does not form any strong side chain interactions with other residues (polar or non-polar). This leaves N211 “free to mutate”, although the strongly stabilising effect of the N211A mutation was not predictable *a priori* on this basis. Another consideration is asparagine has a relatively low helical propensity so switching to alanine may impact on the stability of helix 7 and thus the protein as a whole.

In terms of the impact on substrate specificity, the N211A mutation alone increases activity towards C4 substrates but at the expense of substrate affinity. Structural analysis revealed that M187 plays arguably the main role in forming the plug (Figure 4a). Crucially though the N211A mutation now allows the M187A mutation to be tolerated but with an apparent effect on structural stability based on the CD spectra of the double mutant compared to the N221A single mutant (Figure 5a). The result is that EstN7^DM^ can now catalyse the hydrolysis of C8 substrates, albeit poorly. Indeed, EstN7^DM^ could be considered a more generalist enzyme due to its ability to hydrolyse substrates of different chain lengths. The role of stabilising or suppressor mutations allowing functionally useful but structurally deleterious mutations to be productively expressed in a particular protein is important to the natural and directed evolutionary process (41–43). Here, we inadvertently discovered a suppressor mutation [N211A] in EstN7 but it clearly shows the impact such mutations can have.

In the context of EstN7, W206 appears to play a key role in activity as mutation to alanine results in disruption of both substrate affinity and turnover (Table 1) but does not result in total loss of structural stability as with M187A. While W206 can vary between leucine and phenylalanine in its nearest structural homologues, large hydrophobic amino may be a requisite rather than a small side chain amino acid; W206 is an important hydrophobic tethering residue with one face of the indole ring available to potentially interact with substrate. It could also impose a geometrical restraint on the substrate size and route for the correct approach into the active site, blocking off non-productive access.

The presence of the buried water molecules on the other side of plug does raise the issue of their contribution to the structure-function relationship and how such esterases can be engineered to accept longer chain substrates. While opening the plug has an effect, substrate exchange with water molecules at the end of tunnel (assuming that the mutants fold to hold water in this cavity) and the subsequent polar nature of the cavity may further hinder substrate binding of largely hydrophobic hydrocarbon chains used here. This is one of the potential reasons for the relatively small improvement in activity of EstN7^DM^ towards C4 and longer substrates. Another is conformational and dynamic changes the protein may make to account for the mutations. Thus, extensive secondary engineering efforts will be needed to optimise esterases like EstN7 for use with longer acyl chain compounds. However, the polar nature of expanding the acyl binding pocket into the water cavity may open the opportunity for more specific uses in which the nature of the acyl component is polar.

## Conclusion

Cold-active esterases are important for various industrial and biotechnological processes but limitations on their substrate specificity and range can be a major limiting factor. Using data mining we find that triacetin is potentially a commercially useful substrate to generate a defined single terminally deacylated product. There is an argument that cold-adapted esterases may be less tolerant to protein engineering approaches to expand substrate specificity due to the perceived inherent low thermal stability and the potentially destabilising effect opening up the plug has on esterase structure. EstN7 has relatively high thermal stability, which can be enhanced through a stabilising suppressor mutation (N211A). This in turn allows the main plug opening mutation, M187A to be incorporated. EstN7, in common with related esterases, also has a water-filled cavity beyond the acyl binding pocket plug. This polar, buried cavity may prove to be hinderance in terms of classical screening of engineered variants using substrates like those that we used here. However, it might provide other opportunities for use with more polar substrates. The inherent activity of EstN7 is unknown but analysis of the active site and enzyme kinetics suggests that short chain substrates are likely its target. Thus tunnel geometry, including the shape, polar character and size, is a key determinant for molecular interactions between the ligand and the catalytic site (44).

## Experimental procedures

### Structural analysis

The structure of EstN7^WT^ was determined previously (20). Docking the pNP-C2 and pNP-C8 substrates with the enzyme was performed manually using previous knowledge of esterase structures and the position of the ethylene glycol moiety located in active site of the present structure to guide the process. K4V, the PDB entity defining the C2 substrate, was used, which was manipulated in COOT(28) and placed at geometrically acceptable distances from the surrounding active site lining. This coarse positioning was refined with REFMAC(29) to regularise the geometry of the model. Three different poses were tried, yielding a preferred result, based on best overlap with the ethylene glycol molecule and charge complementarity with the surrounding side chains. When the preferred docking configuration was extended to the C8 substrate version (constructed with JLIGAND(45)), in conjunction with targeted mutations introduced *in silico*, it proved to be the putatively correct orientation upon geometry regularisation with REFMAC as judged by the lack of steric repulsion by the non-polar side chains, and deeper insertion of the extended acyl group into the pocket by around 3.0Å. Energy interactions resulting from the docking of the putative substrates were calculated with PISA, yielding 2.3 kcal/mol. The entropic effect of replacing the waters in the enclosed vestibule with an aliphatic side chain could not be estimated. The lack of higher energy interaction would be consistent with a wide vestibule able to accommodate a variety of entities without clashes. Thermal stability was determined by circular dichroism (CD) spectroscopy as described previously (20).

### Protein engineering and protein production

EstN7 mutants were generated by inverse PCR using the plasmid encoding the EstN7^WT^ gene as a template. The primers pairs used to introduce each mutation were: M187A (Forward 5′-CCAgctATTGATGATAAAAACAATTCACC-3′; Reverse 5′-ATATAATGGCATTTGGAAGCAAAG-3′), N211A (Forward 5′-TCATGATTTAgctGAAAAAGGTTGGTCTAT-3′; Reverse 5′-TTCCAGATTAGATTGCCTGTAATCTC-3′) and W206A (Forward 5′-CTAATCgcGAATCATGATTTAAACG; Reverse 5′-ATTGCCTGTAATCTCTAAGCTG). After verification of the mutations by DNA sequencing, the mutant constructs were used to transform *E. coli* BL21 DE3 cells and the variants produced and purified as previously (20).

### Enzyme activity and kinetics

The protein concentration of purified enzymes was determined using Bradford assay (46). The hydrolytic activity of EstN7 and its variants was determined spectrophotometrically using pNP-derivatives through measuring the liberated *para*-nitrophenol at 410 nm. The reaction was performed at 25°C in 50 mM Tris-HCl buffer (pH 8.0) with enzyme concentration adjusted appropriately enzyme (range 0.005-14μM) depending on rate for a particular substrate. The substrate specificity of the wild-type enzyme and the mutants were determined using different chain length of pNP derivatives; pNP acetate (C2), pNP butyrate (C4), pNP hexanpate (C6) and pNP octanoate (C8). Each substrate was prepared as a 10mM stock in dimethyl dulfoxide (DMSO). The kinetic parameters, Michaelis constant (*K*_M_) and maximum rate (V_max_) and turnover number (*k*_cat_) were calculated using Hyper32 (https://hyper32.software.informer.com/).

EstN7 hydrolytic activity towards triacetin as a substrate was assessed by pH end point titration. The reaction was performed at 20°C in a glass vessel containing 15mL Tris–HCl buffer (50mM), 270 mM triacetin, and suitable enzyme concentration (0.5 mg/ml final concentration). At the end of the incubation period, the liberated free fatty acids were titrated against 0.03M NaOH using phenolphthalein as an indicator. A control sample was tested in parallel to determine the original free fatty acid content in the substrate. Product was also confirmed by NMR as outlined in the Supporting Information.

### Data-mining for new valuable biotransformations

A bespoke version of the enzyme biotransformation data-mining software Transform-MinER (47) was created using experimentally confirmed substrates and products of reactions catalysed by EstN7. This bespoke version will be referred to as the EstN7 version of Transform-MinER. A public web-server (31) based on reactions in the KEGG database (48) has been made available but a bespoke version enables targeted search against the confirmed reactions. Virtual screening of a dataset of ~400,000 commercially available molecules was performed using the EstN7 version of Transform-MinER, identifying reactions with high similarity to known EstN7 reactions. These were subsequently assessed for commercial interest by obtaining recent purchase prices for substrates and products from online sources and filtering for reactions with good margins. EstN7-triacetin docking was performed using HADDOCK 2.4 web server (https://wenmr.science.uu.nl/haddock2.4/) (49) with the coordinates of triacetin obtained from PubChem (https://pubchem.ncbi.nlm.nih.gov) with the PubChem CID 5541.

## Data availability

The structure of EstN7 has been deposited in the PDB under accession code 7b4q. All other data is available on request from the authors.

## Supporting Information

This article contains supporting information: Supporting Table 1, Supporting Figure S1-S5.

## Acknowledgements

We would like to thank the Protein Technology Hub facility in the School of Biosciences, Cardiff University for access to protein purification and analysis facilities. We would like to thank Dr.Mohamed Teleb, Lecturer of Organic chemistry, faculty of Pharmacy, Alexandria University, for his help in NMR data analysis. A special thank you to Prof. Ahmed Rafik El-Mahdy, professor of Biochemistry, faculty of Agriculture, Alexandria University for his guidance.

## References

1. Prakash, D., Nawani, N., Prakash, M., Bodas, M., Mandal, A., Khetmalas, M., and Kapadnis, B. (2013) Actinomycetes: a repertory of green catalysts with a potential revenue resource. BioMed research international 2013

2. Sheldon, R. A. (2016) Engineering a more sustainable world through catalysis and green chemistry. Journal of The Royal Society Interface 13, 20160087

3. Wang, M., Si, T., and Zhao, H. (2012) Biocatalyst development by directed evolution. Bioresource technology 115, 117–125

4. Dunn, P. J. (2012) The importance of green chemistry in process research and development. Chemical Society Reviews 41, 1452–1461

5. Lorenz, P., and Eck, J. (2005) Metagenomics and industrial applications. Nat Rev Microbiol 3, 510–516

6. Bornscheuer, U. T. (2002) Microbial carboxyl esterases: classification, properties and application in biocatalysis. FEMS Microbiol Rev 26, 73–81

7. Jaeger, K. E., Dijkstra, B. W., and Reetz, M. T. (1999) Bacterial biocatalysts: molecular biology, three-dimensional structures, and biotechnological applications of lipases. Annu Rev Microbiol 53, 315–351

8. Chandra, P., Enespa, Singh, R., and Arora, P. K. (2020) Microbial lipases and their industrial applications: a comprehensive review. Microb Cell Fact 19, 169

9. Sarmiento, F., Peralta, R., and Blamey, J. M. (2015) Cold and hot extremozymes: industrial relevance and current trends. Frontiers in bioengineering and biotechnology 3, 148

10. Al-Ghanayem, A. A., and Joseph, B. (2020) Current prospective in using cold-active enzymes as eco-friendly detergent additive. Applied Microbiology and Biotechnology 104, 2871–2882

11. Al-Maqtari, Q. A., Waleed, A.-A., and Mahdi, A. A. (2019) Cold-active enzymes and their applications in industrial fields-A review.

12. Noby, N., Hussein, A., Saeed, H., and Embaby, A. M. (2020) “Recombinant cold-adapted halotolerant, organic solvent-stable esterase (estHIJ) from Bacillus halodurans. Analytical Biochemistry 591, 113554

13. Joseph, B., Ramteke, P. W., and Thomas, G. (2008) Cold active microbial lipases: some hot issues and recent developments. Biotechnology advances 26, 457–470

14. Kuddus, M. (2015) Cold-active microbial enzymes. Biochem Physiol 4, e132

15. Elend, C., Schmeisser, C., Hoebenreich, H., Steele, H., and Streit, W. (2007) Isolation and characterization of a metagenome-derived and cold-active lipase with high stereospecificity for (R)-ibuprofen esters. Journal of biotechnology 130, 370–377

16. Rotticci, D., Ottosson, J., Norin, T., and Hult, K. (2001) Candida antarctica Lipase BA Tool for the Preparation of Optically Active Alcohols. in Enzymes in Nonaqueous Solvents, Springer. pp 261–276

17. Maharana, A., and Ray, P. (2015) A novel cold-active lipase from psychrotolerant Pseudomonas sp. AKM-L5 showed organic solvent resistant and suitable for detergent formulation. Journal of Molecular Catalysis B: Enzymatic 120, 173–178

18. Santiago, M., Ramírez-Sarmiento, C. A., Zamora, R. A., and Parra, L. P. (2016) Discovery, molecular mechanisms, and industrial applications of cold-active enzymes. Frontiers in microbiology 7, 1408

19. Jemli, S., Ayadi-Zouari, D., Hlima, H. B., and Bejar, S. (2016) Biocatalysts: application and engineering for industrial purposes. Critical reviews in biotechnology 36, 246–258

20. Noby, N., and Jones, D. (2021) TBC. TBC TBC, TBC

21. Noby, N., Saeed, H., Embaby, A. M., Pavlidis, I. V., and Hussein, A. (2018) Cloning, expression and characterization of cold active esterase (EstN7) from Bacillus cohnii strain N1: A novel member of family IV. International journal of biological macromolecules 120, 1247–1255

22. Chovancova, E., Pavelka, A., Benes, P., Strnad, O., Brezovsky, J., Kozlikova, B., Gora, A., Sustr, V., Klvana, M., Medek, P., Biedermannova, L., Sochor, J., and Damborsky, J. (2012) CAVER 3.0: a tool for the analysis of transport pathways in dynamic protein structures. PLoS Comput Biol 8, e1002708

23. Tutino, M. L., di Prisco, G., Marino, G., and de Pascale, D. (2009) Cold-adapted esterases and lipases: from fundamentals to application. Protein Pept Lett 16, 1172–1180

24. Levisson, M., Han, G. W., Deller, M. C., Xu, Q., Biely, P., Hendriks, S., Ten Eyck, L. F., Flensburg, C., Roversi, P., Miller, M. D., McMullan, D., von Delft, F., Kreusch, A., Deacon, A. M., van der Oost, J., Lesley, S. A., Elsliger, M. A., Kengen, S. W., and Wilson, I. A. (2012) Functional and structural characterization of a thermostable acetyl esterase from Thermotoga maritima. Proteins 80, 1545–1559

25. McKary, M. G., Abendroth, J., Edwards, T. E., and Johnson, R. J. (2016) Structural Basis for the Strict Substrate Selectivity of the Mycobacterial Hydrolase LipW. Biochemistry 55, 7099–7111

26. Zhu, X., Larsen, N. A., Basran, A., Bruce, N. C., and Wilson, I. A. (2003) Observation of an arsenic adduct in an acetyl esterase crystal structure. J Biol Chem 278, 2008–2014

27. Palm, G. J., Fernandez-Alvaro, E., Bogdanovic, X., Bartsch, S., Sczodrok, J., Singh, R. K., Bottcher, D., Atomi, H., Bornscheuer, U. T., and Hinrichs, W. (2011) The crystal structure of an esterase from the hyperthermophilic microorganism Pyrobaculum calidifontis VA1 explains its enantioselectivity. Appl Microbiol Biotechnol 91, 1061–1072

28. Emsley, P., and Cowtan, K. (2004) Coot: model-building tools for molecular graphics. Acta Crystallogr D Biol Crystallogr 60, 2126–2132

29. Murshudov, G. N., Skubak, P., Lebedev, A. A., Pannu, N. S., Steiner, R. A., Nicholls, R. A., Winn, M. D., Long, F., and Vagin, A. A. (2011) REFMAC5 for the refinement of macromolecular crystal structures. Acta Crystallogr D Biol Crystallogr 67, 355–367

30. Hotta, Y., Ezaki, S., Atomi, H., and Imanaka, T. (2002) Extremely stable and versatile carboxylesterase from a hyperthermophilic archaeon. Appl Environ Microbiol 68, 3925–3931

31. Tyzack, J. D., Ribeiro, A. J. M., Borkakoti, N., and Thornton, J. M. (2018) Transform-MinER: transforming molecules in enzyme reactions. Bioinformatics 34, 3597–3599

32. Chen, T., Gong, T., Zhao, T., Liu, X., Fu, Y., Zhang, Z., and Gong, T. (2017) Paclitaxel loaded phospholipid-based gel as a drug delivery system for local treatment of glioma. Int J Pharm 528, 127–132

33. Feller, G., and Gerday, C. (2003) Psychrophilic enzymes: hot topics in cold adaptation. Nat Rev Microbiol 1, 200–208

34. Liu, Y., Xu, H., Yan, Q., Yang, S., Duan, X., and Jiang, Z. (2013) Biochemical characterization of a first fungal esterase from Rhizomucor miehei showing high efficiency of ester synthesis. PLoS One 8, e77856

35. Sugihara, A., Shimada, Y., Nomura, A., Terai, T., Imayasu, M., Nagai, Y., Nagao, T., Watanabe, Y., and Tominaga, Y. (2002) Purification and characterization of a novel cholesterol esterase from Pseudomonas aeruginosa, with its application to cleaning lipid-stained contact lenses. Bioscience, biotechnology, and biochemistry 66, 2347–2355

36. Komiya, D., Hori, A., Ishida, T., Igarashi, K., Samejima, M., Koseki, T., and Fushinobu, S. (2017) Crystal Structure and Substrate Specificity Modification of Acetyl Xylan Esterase from *Aspergillus luchuensis*. Applied and Environmental Microbiology 83, e01251–01217

37. Juhl, P. B., Doderer, K., Hollmann, F., Thum, O., and Pleiss, J. (2010) Engineering of Candida antarctica lipase B for hydrolysis of bulky carboxylic acid esters. Journal of biotechnology 150, 474–480

38. Santarossa, G., Lafranconi, P. G., Alquati, C., DeGioia, L., Alberghina, L., Fantucci, P., and Lotti, M. (2005) Mutations in the “lid” region affect chain length specificity and thermostability of a Pseudomonas fragi lipase. FEBS letters 579, 2383–2386

39. Holmquist, M. (1998) Insights into the molecular basis for fatty acyl specificities of lipases from Geotrichum candidum and Candida rugosa. Chemistry and physics of lipids 93, 57–65

40. Pereira, M. R., Maester, T. C., Mercaldi, G. F., de Macedo Lemos, E. G., Hyvönen, M., and Balan, A. (2017) From a metagenomic source to a high-resolution structure of a novel alkaline esterase. Applied microbiology and biotechnology 101, 4935–4949

41. Peisajovich, S. G., and Tawfik, D. S. (2007) Protein engineers turned evolutionists. Nat Methods 4, 991–994

42. Bershtein, S., Segal, M., Bekerman, R., Tokuriki, N., and Tawfik, D. S. (2006) Robustness-epistasis link shapes the fitness landscape of a randomly drifting protein. Nature 444, 929–932

43. Sideraki, V., Huang, W., Palzkill, T., and Gilbert, H. F. (2001) A secondary drug resistance mutation of TEM-1 beta-lactamase that suppresses misfolding and aggregation. Proc Natl Acad Sci U S A 98, 283–288

44. Kingsley, L. J., and Lill, M. A. (2015) Substrate tunnels in enzymes: structure– function relationships and computational methodology. Proteins: Structure, Function, and Bioinformatics 83, 599–611

45. Lebedev, A. A., Young, P., Isupov, M. N., Moroz, O. V., Vagin, A. A., and Murshudov, G. N. (2012) JLigand: a graphical tool for the CCP4 template-restraint library. Acta Crystallographica Section D 68, 431–440

46. Bradford, M. M. (1976) A rapid and sensitive method for the quantitation of microgram quantities of protein utilizing the principle of protein-dye binding. Analytical biochemistry 72, 248–254

47. Tyzack, J. D., Ribeiro, A. J. M., Borkakoti, N., and Thornton, J. M. (2019) Exploring Chemical Biosynthetic Design Space with Transform-MinER. ACS Synth Biol 8, 2494–2506

48. Kanehisa, M., Sato, Y., Kawashima, M., Furumichi, M., and Tanabe, M. (2016) KEGG as a reference resource for gene and protein annotation. Nucleic Acids Res 44, D457–462

49. van Zundert, G. C. P., Rodrigues, J., Trellet, M., Schmitz, C., Kastritis, P. L., Karaca, E., Melquiond, A. S. J., van Dijk, M., de Vries, S. J., and Bonvin, A. (2016) The HADDOCK2.2 Web Server: User-Friendly Integrative Modeling of Biomolecular Complexes. J Mol Biol 428, 720–725

