## Supplementary material for "Structure guided engineering of a cold active esterase expands substrate range though a stabilisation mutation that allows access to a buried water chamber": SI

**Supporting information.**

**Figure S1.**

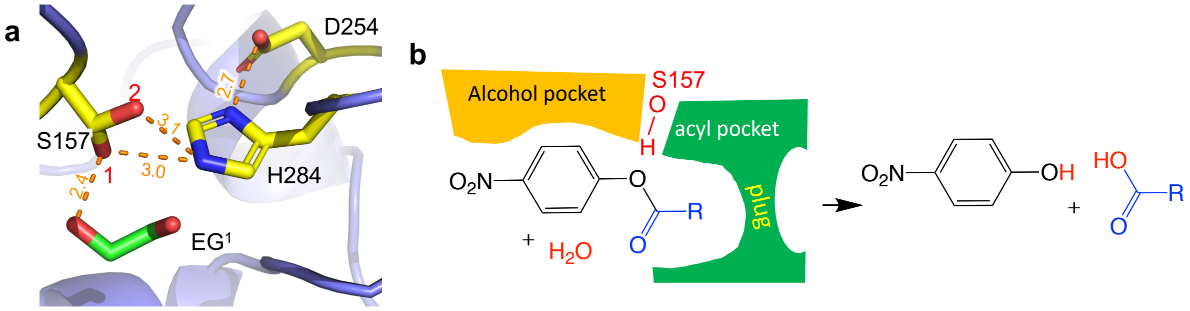


Hydrolysis of pNA substrate. (a) Close up for the active site with the catalytic triad residues coloured yellow. S157 has two rotamers labelled 1 and 2. Orange dashed lines represented the distances between individual atoms. $EG^1$ is an ethylene glycol molecule found in the active site. (b)The esterase alcohol and acyl binding pockets, including the acyl pocket plug are shown schematically as are the products of the reaction involving the pNP substrate. The R group represents different linear alkane side chain lengths mentioned in the main manuscript.

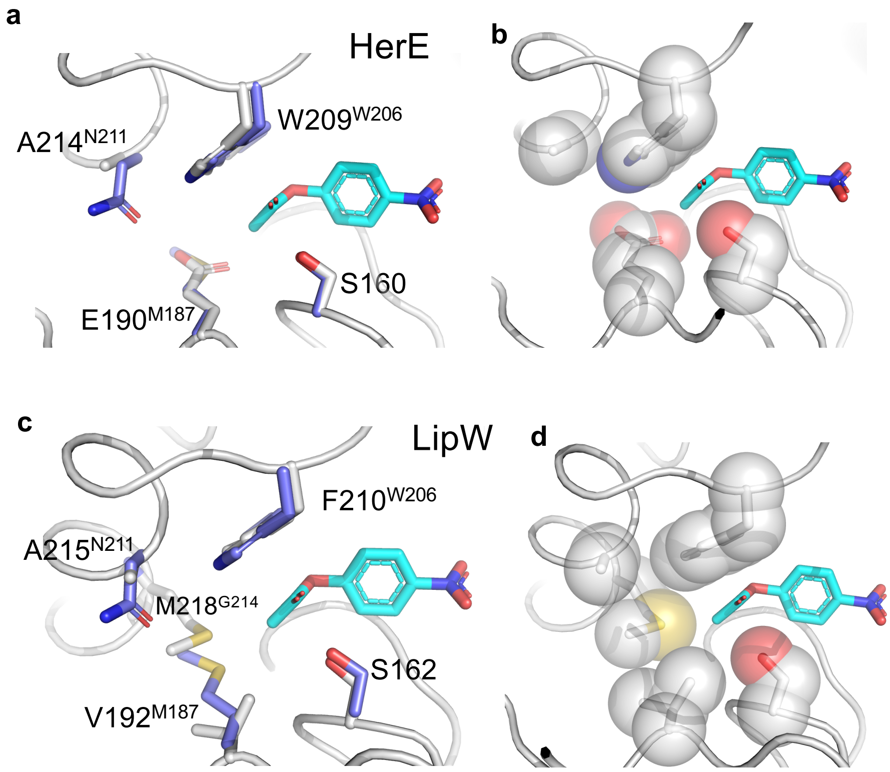


**Figure S2.** Comparison of EstN7 the acyl binding pocket with HerE (a,c) and LipW (1) (b,d) (PDB 3QH4). The pNP-C2 substrate is shown in cyan. Panels a and c show the structural overlap of EstN7 (blue) with its counterpart (grey). Panels c and d show the key residues involved in forming the acyl pocket plus for HerE and LipW, respectively.

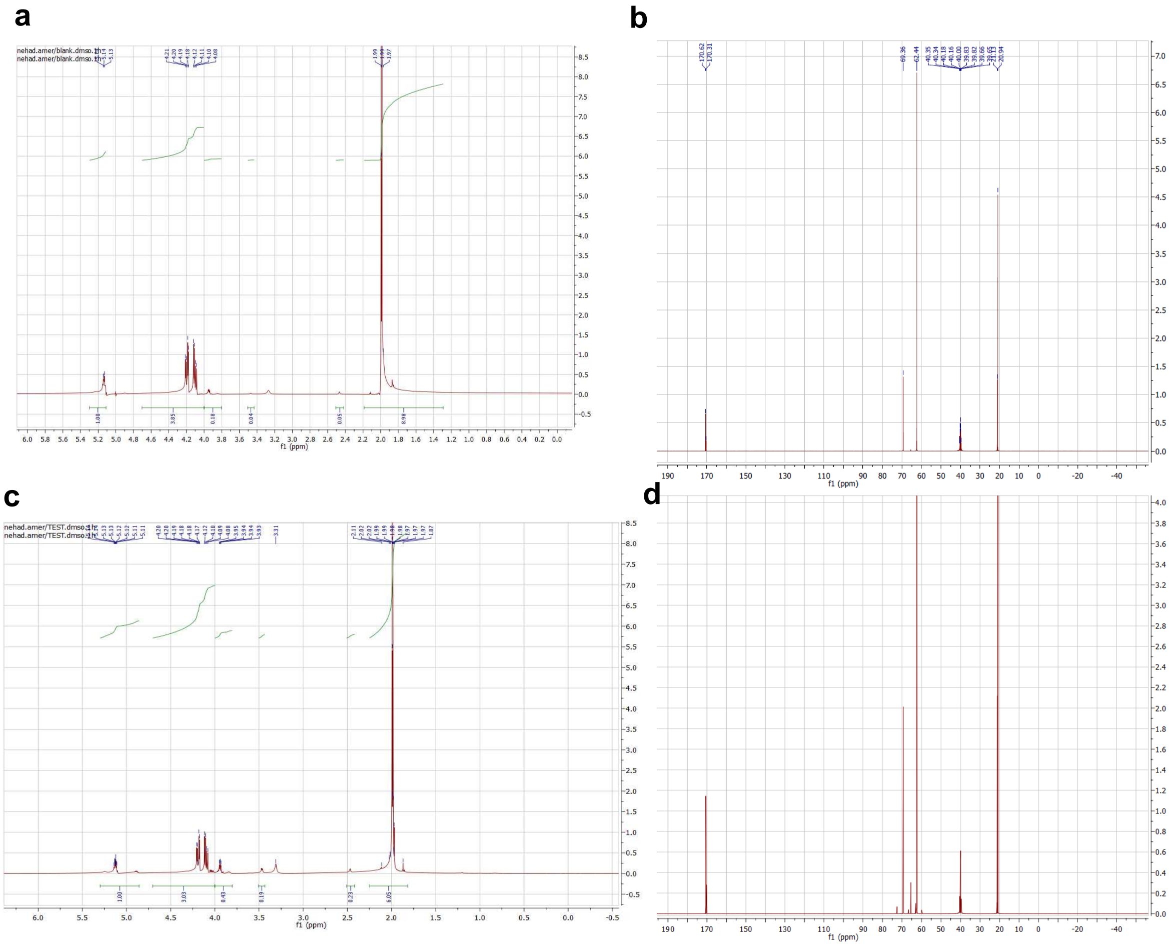


**Figure S3. NMR analysis of triacetin and it breakdown product after incubation with EstN7.** (a-b) The $^{1}H$ (a) and $^{13}C$ 1D NMR spectra of triacetin. (c-d) The $^{1}H$ (c) and (d) $^{13}C$ 1D NMR spectra of reaction products. Nuclear magnetic resonance spectra ($^{1}H$-NMR and $^{13}C$-NMR) were recorded on JEOL 500 MHz spectrometers at ambient temperature operating at 500 and 125 MHz, respectively. Chemical shifts were reported in parts per million (ppm) and are referenced relative to DMSO at δ 2.50 ppm for DMSO-$d_6$.

**Method**. The enzyme regioselectivity towards triacetin was determined by NMR. The hydrolysis reaction was performed as stated previously, and at the end of the incubation period, the enzyme was removed from the reaction using affinity chromatography. 3 volumes of ethyl acetate were added for liquid portioning process to extract the hydrolytic products, the extraction was repeated at least 3 times. The collected ethyl acetate was totally evaporated and the remaining products were analysed by NMR using DMSO as a solvent. A blank sample was typically processed as described above.

**Spectral Analysis.** The overall structure of triacetin is shown in Figure 5a of the main manuscript, including the annotation of the central glyceryl unit. The $^{1}H$-NMR spectrum of triacetin (Figure S3a) shows a singlet peak at 1.99 ppm chemical shift equivalent to 9 protons, which corresponds to the three methyl groups; a multiplet peak is observed at the range 4.08-4.21 ppm equivalent to 4 protons corresponding to the two (C-1 and C-3 of the glyceryl unit) methylene protons. The multiplet peak at the range 5.12-5.15 ppm equivalent to one proton correspond to the CH (C-2 of the glyceryl unit) proton. In addition, the $^{13}C$-NMR spectrum of triacetin (Figure S3b) shows 5 peaks at 20.94, 62.44, 69.36, 170.31, 170.62 ppm, corresponding to $CH_3$, $CH_2$, CH, C=O, C=O groups.

On the other hand, the $^{1}$H-NMR spectrum of the reaction product (Figure S3c), shows additional multiplet peaks, than those observed for triacetin at the range 3.44-3.49, 3.70-3.98, and 4.6-4.9 ppm. In addition, the integration of the peak at 1.99 ppm corresponding to the methyl groups, is equivalent to 6 protons, confirming the hydrolysis of one methyl group. The $^{13}$C-NMR spectrum of the reaction product (Figure S3d), also shows a slightly deshielded peak at 65 ppm corresponding to either C-1 or C-3 of the glyceryl unit, which underwent hydrolysis. The hydrolysis at the central C-2 carbon of the glyceryl will give a symmetric reaction product resulting in significant deshielding of the peak corresponding to the C-2-H proton, which is not observed in the $^{13}$C NMR spectra of the product (Figure S3d). The observed results confirm that hydrolysis occurred at either the terminal 1 or 3 carbon position.

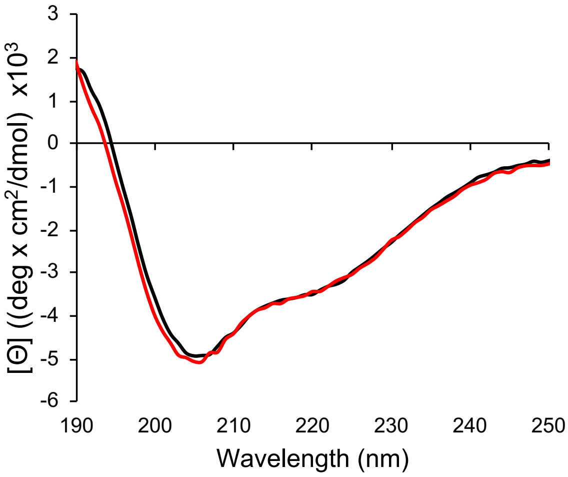


**Figure S4.** CD spectra of EstN7$^{WT}$ (black) and EstN7$^{N211A}$ (red) were measured at 70°C after temperature ramping.

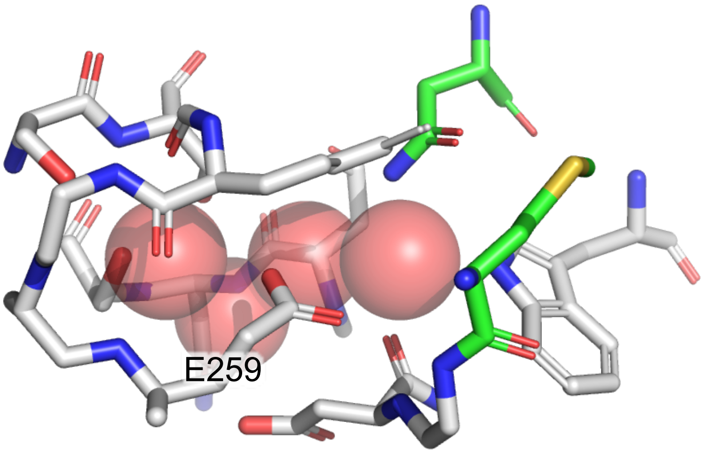


**Figure S5.** Residues surrounding the water molecules in the second cavity beyond the plug. Residues coloured green are N211 and M187 that comprise the plug. E259 is highlighted as it is the only residue capable of making an ionic interaction with the water molecules.

### Supporting References


1. McKary, M. G., Abendroth, J., Edwards, T. E., and Johnson, R. J. (2016) Structural Basis for the Strict Substrate Selectivity of the Mycobacterial Hydrolase LipW. *Biochemistry* **55**, 7099-7111
